# Oxford Nanopore sequencing-based protocol to detect CpG methylation in human mitochondrial DNA

**DOI:** 10.1101/2021.02.20.432086

**Authors:** Iacopo Bicci, Claudia Calabrese, Zoe J. Golder, Aurora Gomez-Duran, Patrick F Chinnery

## Abstract

Methylation on CpG residues is one of the most important epigenetic modifications of nuclear DNA, regulating gene expression. Methylation of mitochondrial DNA (mtDNA) has been studied using whole genome bisulfite sequencing (WGBS), but recent evidence has uncovered major technical issues which introduce a potential bias during methylation quantification. Here, we validate the technical concerns with WGBS, and then develop and assess the accuracy of a protocol for variant-specific methylation identification using long-read Oxford Nanopore Sequencing. Our approach circumvents mtDNA-specific confounders, while enriching for native full-length molecules over nuclear DNA. Variant calling analysis against Illumina deep re-sequencing showed that all expected mtDNA variants can be reliably identified. Methylation calling revealed negligible mtDNA methylation levels in multiple human primary and cancer cell lines. In conclusion, our protocol enables the reliable analysis of epigenetic modifications of mtDNA at single-molecule level at single base resolution, with potential applications beyond methylation.

**Motivation:** Although whole genome bisulfite sequencing (WGBS) is the gold-standard approach to determine base-level CpG methylation in the nuclear genome, emerging technical issues raise questions about its reliability for evaluating mitochondrial DNA (mtDNA) methylation. Concerns include mtDNA strand asymmetry rendering the C-rich light strand disproportionately vulnerable the chemical modifications introduced with WGBS. Also, short-read sequencing can result in a co-amplification of nuclear sequences originating from ancestral mtDNA with a high nucleotide similarity. Lastly, calling mtDNA alleles with varying proportions (heteroplasmy) is complicated by the C-to-T conversion introduced by WGBS on unmethylated CpGs. Here, we propose an alternative protocol to quantify methyl-CpGs in mtDNA, at single-molecule level, using Oxford Nanopore Sequencing (ONS). By optimizing the standard ONS library preparation, we achieved selective enrichment of native mtDNA and accurate single nucleotide variant and CpG methylation calling, thus overcoming previous limitations.

## Introduction

Cytosine methylation is an epigenetic modification of nuclear DNA (nDNA) that can regulate gene expression during development (Smith and Meissner, 2013) and throughout life (Siegfried and Simon, 2010), but the presence of CpG methylation of the mitochondrial genome (mtDNA) is a matter of debate. This is an important issue to resolve given the pivotal role of mtDNA in cellular metabolism (Suomalainen and Battersby, 2018).

Whole genome bisulfite sequencing (WGBS) is the gold standard technique for detecting methylation across the nuclear genome (nDNA) (Krueger *et al*., 2012; Wolters *et al*., 2017; Sirard, 2019), where sequencing before and after the chemical conversion of unmethylated cytosine to uracil allows the degree of methylation to be measured at single-base resolution. WGBS studies have reported methylation patterns across the mtDNA molecule in different biological contexts (Devall *et al*., 2017). However, recent studies suggest that these are influenced by technical artefacts (Hong *et al*., 2013; Liu *et al*., 2016; Mechta *et al*., 2017). MtDNA has a purine-rich “Heavy”(H-) and a pyrimidine-rich “Light”(L-) strand (Anderson *et al*., 1981), leading to a disproportionate fragmentation of the cytosine-rich L-strand by bisulfite treatment (Olova *et al*., 2018). Moreover, the presence of multiple mtDNA genotypes within mitochondria of the same cell (heteroplasmy (Stewart and Chinnery, 2020)), and nuclear sequences originated from the mtDNA (NUMTs (Hazkani-Covo, Zeller and Martin, 2010; Dayama *et al*., 2014)) are potential confounders for mtDNA methylation detection.

To overcome these limitations we set out to quantify CpG methylation of native mtDNA using long-read based Oxford Nanopore Sequencing (ONS) technology (Jain *et al*., 2016). The core of our protocol is an enzymatic digestion of genomic DNA (gDNA) followed by selective enrichment of the longer fragments, in order to retain linearised, full-length native mtDNA molecules in this fraction. Given recent evidence that ONS can be used to measure differentially methylated nuclear imprinted genes (Gigante *et al*., 2019), and that long-read sequencing enables the detection of NUMTs breakpoints in the nucleus (Wei *et al*., 2020), our approach represent an improvement of the standard ONS library preparation protocol based on random fragmentation of the gDNA. We further improve methylation detection by studying the accuracy of the methylation calling on mtDNA, using negative and positive controls, while simultaneously assessing the validity of the variant calling by comparing our results with Illumina sequencing.

Finally, we investigated the presence of base-level mtDNA methylation in human cancer and primary cell lines, showing negligible levels of mtDNA CpG methylation.

## Results

### CpG methylation analysis of mtDNA with WGBS

We sought independent evidence that WGBS has limitations for mtDNA by analysing data from 67 human cell lines and tissues from the NIH Human Epigenome Roadmap Project (Roadmap Epigenomics Consortium *et al*., 2015). Fifty-five passed quality control (Methods) and were aligned to the human genome build GRCh38 (**Data S1)**. Analysis of the mtDNA-aligned reads revealed a pronounced *per*-strand mapping and coverage bias observed in 58.2% (N= 32) samples (here termed the “Biased” group, BG, **Figure 1A-B, Figure S1A**). The majority of reads in the BG mapped to the mitochondrial H-strand (≥ 55% reads; *P* = ≤ 0.0001, **Figure 1A, Figure S1A**) and samples showed a more pronounced *per*-strand coverage bias on the L-strand (L-strand coverage_BG_ = 6.2%-88.3%; H-strand coverage_BG_ = 83.5%-91.7%, **Figure 1B** top panel). The remaining data (N = 23, “Low Bias” group), showed a milder mapping bias on the H-strand (between 51%-55% reads; *P* = ≤ 0.0001, **Figure 1A, Figure S1A**) but no coverage bias (**Figure 1B** bottom panel). We observed differences (Mann-Whitney test: *P* = ≤ 0.0001) in the average read depth per position calculated in the two groups: 66.32 ± 28.84x BG versus 148.77 ± 55.45x LBG (mean ± sd; **Figure 1C**). We hence performed methylation analysis in both groups and found higher apparent methylation levels in the L-strand compared to the H in all samples analyzed (L-strand_BG_= 4.97% ± 8.79 vs H-strand_BG_= 2.01% ± 1.92 (mean methylation ± sd); L-strand_LBG_= 1.43% ± 0.77 vs H-strand_LBG_= 1.39% ± 0.7 (mean methylation ± sd); *P* = ≤ 0.001; **Figure 1D, Figure S1B**). This is explained by a significant inverse correlation with the read depth per position (Spearman”s rank test P < 2.2e-16; average rho coefficient = −0.78, **Figure 1E**), leading to the appearance of higher methylation levels where read depth is low. This holds true also for the Low Bias group (with a milder alignment bias), where local fluctuations in the read depth alter CpG methylation levels (**Figure S1C-D**). This is consistent with previous observations indicating a bisulfite-related selective loss of the cytosine-rich L-strand (Olova *et al*., 2018).

**Figure 1.**
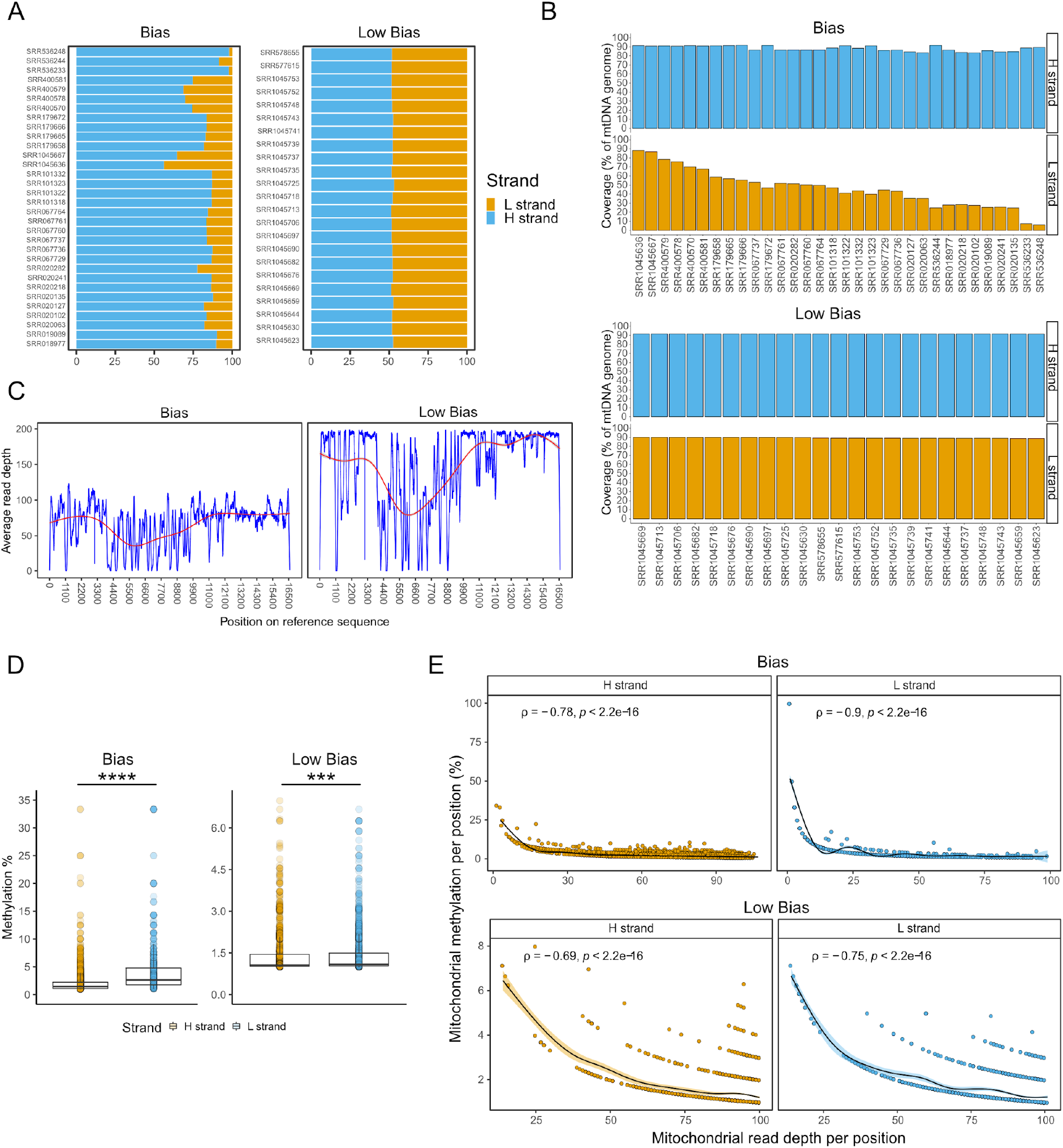
Quantification of WGBS alignment and coverage bias. **a)** Percentage of reads aligned to the mtDNA reference per sample, identifying samples with a marked (Bias, N = 32) or low (Low Bias, N = 23) per-strand-bias. **b)** Percentage of mtDNA covered by at least 5 reads on the two mtDNA strands (H and L) in (top) Bias and (bottom) Low Bias sample groups. **c)** Distribution of the average read depth per mtDNA position in the two per-strand-bias groups. The red lines indicate the mean over all the data points (calculated using the “loess” *geom_smooth* R function). **d**) Quantification of the average CpG methylation per strand (H and L), divided by per-strand-bias group. The lower and upper hinges of the boxplot correspond to the first and third quartile of the distribution, with median in the center and whiskers span no further than 1.5*interquartile range. Stars indicate significance (***: two-sided P ≤ 0.001; two-sided ****: P ≤ 0.0001, Wilcoxon test). **e**) Correlations between average read depth and average methylation percentage for every cytosine in CpG context, in the two groups of per-strand-bias groups and mtDNA strands (H and L). Pearson correlation coefficient and two-sided P-values are shown. For all the plots in **d-e)**, Average methylation is intended as the mean methylation value across all the WGBS samples analyzed.

### Design and assessment of an ONS-based protocol for mtDNA CpG methylation analysis

To overcome the problems intrinsic to the WGBS methylation determination, we set out to quantify mtDNA CpG methylation using ONS on native human genomic DNA. We first established the accuracy of the methylation calling on mtDNA by sequencing a near complete PCR amplicon of human mtDNA (negative control, NC, 0% methylated) and a corresponding positive control generated *in vitro* with a recombinant CpG methyltransferase (= PC, 100% methylated) (**Figure S2A-B**). We used Nanopolish software (Simpson *et al*., 2017) to call methylation on PC and NC, which generated log-likelihood ratio (LLR) values of CpG methylation (**Figure 2A)**. A site is considered methylated when its LLR is above a certain threshold. To choose the most accurate methylation calling threshold for mtDNA, we determined the ratios of true and false positives by varying LLR thresholds values, following previous procedures (Simpson *et al*., 2017) (Methods). We therefore calculated a receiving operating characteristic (ROC) and methylation calling accuracy (intended as proportion of true calls) (**Figure 2B-C**). The ability to distinguish between mtDNA unmethylated and methylated sites was measured by area under the ROC curve (AUC), which was equal to 0.97 (**Figure 2B**). With the default Nanopolish LLR threshold (≥2.5), an accuracy of 97.7% could be achieved (**Figure 2C**). Hence, we chose a more stringent methylation calling threshold (LLR ≥ 5) yielding an accuracy of 99% (**Figure 2C**). Also, by looking at methylation profiles of NC, we identified 13 false positive methylated residues (**Table S1, Figure S2B**), that were further removed in downstream analyses of biological samples (Methods).

**Figure 2.**
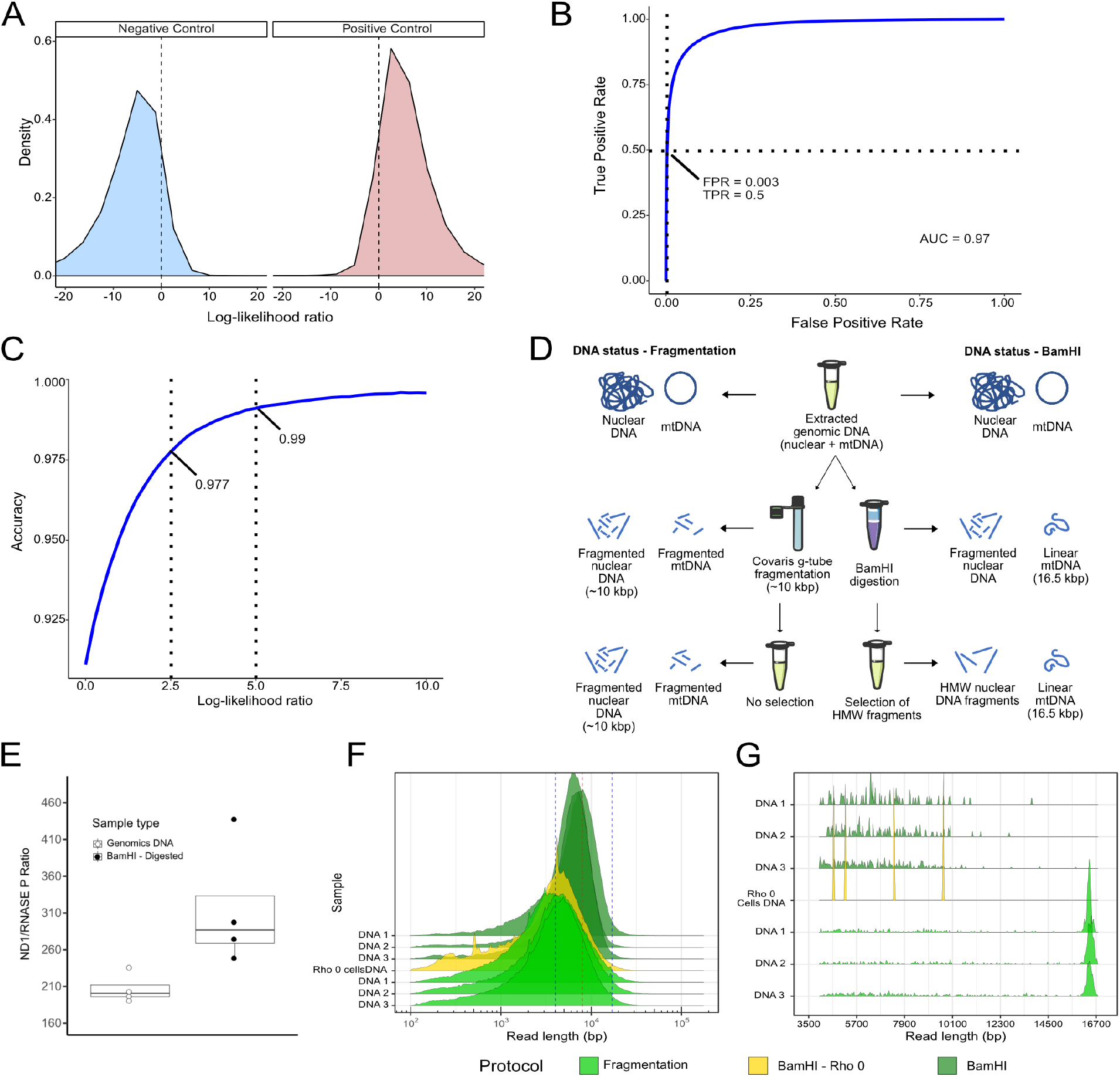
Overview of the proposed ONS-based protocol for mtDNA methylation calling. **a**) Log-likelihood ratio values of methylation calculated by Nanopolish, using the positive and negative controls. The log-likelihood range between −20 and 20 (used to build the ROC curve) is shown. **b**) Receiver operating characteristic (ROC) curve calculated by changing the methylation call log-likelihood ratio threshold from a value of −20 to 20, with a step of 0.25. The dash lines are drawn at FPR (False Positive Rate) and TPR (True Positive Rate) values obtained by setting the ratio equal to 5. AUC = area under the curve. **c**) Methylation call accuracy calculated at increasing values of log-likelihood ratio (ranging between 0 and 10). The dash line indicates the accuracy achieved at the ratio equal to 5 (accuracy = 0.99). **d)** Overview of the workflow used to process samples using (left) standard ONT fragmentation protocol and (right) BamHI-based protocol. **e)** Ratio of signal from the mitochondrial *MT-ND1* over *RNASE P* ddPCR probes in undigested genomic DNA and BamHI-digested genomic DNA. N = 4 for each protocol used. Star indicates significance (*: two-sided *P* = ≤ 0.05, Wilcoxon test). **f)** Distributions of the total sequenced reads in 3 samples prepared with either fragmentation or BamHI-based protocols. Reads from Rho 0 cells (treated with BamHI) are highlighted in yellow. Blue dashed lines correspond to the chosen cutoff for read filtering at 4000bp and 17000 bp. The red dashed line corresponds to the human mtDNA genome length (16.5 Kbp). **g)** Distribution of the aligned reads filtered by length (4000bp −17000 bp) on the human mtDNA reference sequence in 3 samples prepared with either fragmentation or BamHI-based protocols. Reads from Rho 0 cells (treated with BamHI) are highlighted in yellow.

Next, we developed a custom-made library preparation protocol (**Figure 2D**) based on the simultaneous linearisation and enrichment of the native full-length mtDNA molecule (**Figure 2E**) through BamHI restriction enzyme digestion (which usually cuts the mtDNA once). We tested the efficiency of our modified protocol over the standard ONS library preparation, based on random fragmentation, by performing ONS on biological replicates of human DNA (N = 3 different gDNA, 5 technical replicates each, 15 in total). We further performed a strict filtering on read lengths (selecting between 4000 and 17000 bp) and per read quality (Phred ≥ 9) before the alignment (**Figure S3A-B**), followed by supplementary alignments removal. While not altering quality parameters (**Figure S3C-D**), our filtering enriched for full length mtDNA sequences in all BamHI-treated samples and reduced NUMT contamination. This was determined by sequencing rho0 cells lacking mtDNA (King and Attardi, 1989), where we identified only 4 mtDNA-aligned reads with mapping quality of 0 (**Figure 2F-G**).

Under these conditions, the fragmentation-based method showed an L-strand bias (L-strand_FRAG_= 46.12% ± 5.13, H-strand_FRAG_= 53.87% ± 5.13, mean methylation ± sd; Anova one-way test *P* ≤ 0.001, **Figure 3A, Figure S4A**) with 6 samples having < 100% coverage (**Figure 3B**). On the contrary, the BamHI-based protocol did not show any alignment bias (L-strand_BAMHI_ = 50.67% ± 4.07, H-strand_BAMHI_ = 49.32% ± 4.07, mean methylation ± sd; *P* = 0.36, **Figure 3A**) or coverage bias (**Figure 3B**). Average mtDNA read depth was higher in the samples processed with the BamHI-based protocol (Frag. = 23.83x ± 4.33, BamHI = 131.73x ± 8.15, mean ± sd; *P* = ≤ 0.0001, **Figure 3C, Figure S4B**), with almost half of the mitochondrial reads mapped as full-length molecules (≥ 15,000 bp; 42% ± 12 of BamHI reads Vs 2% ± 2 of Frag. reads, **Figure 2G**).

**Figure 3.**
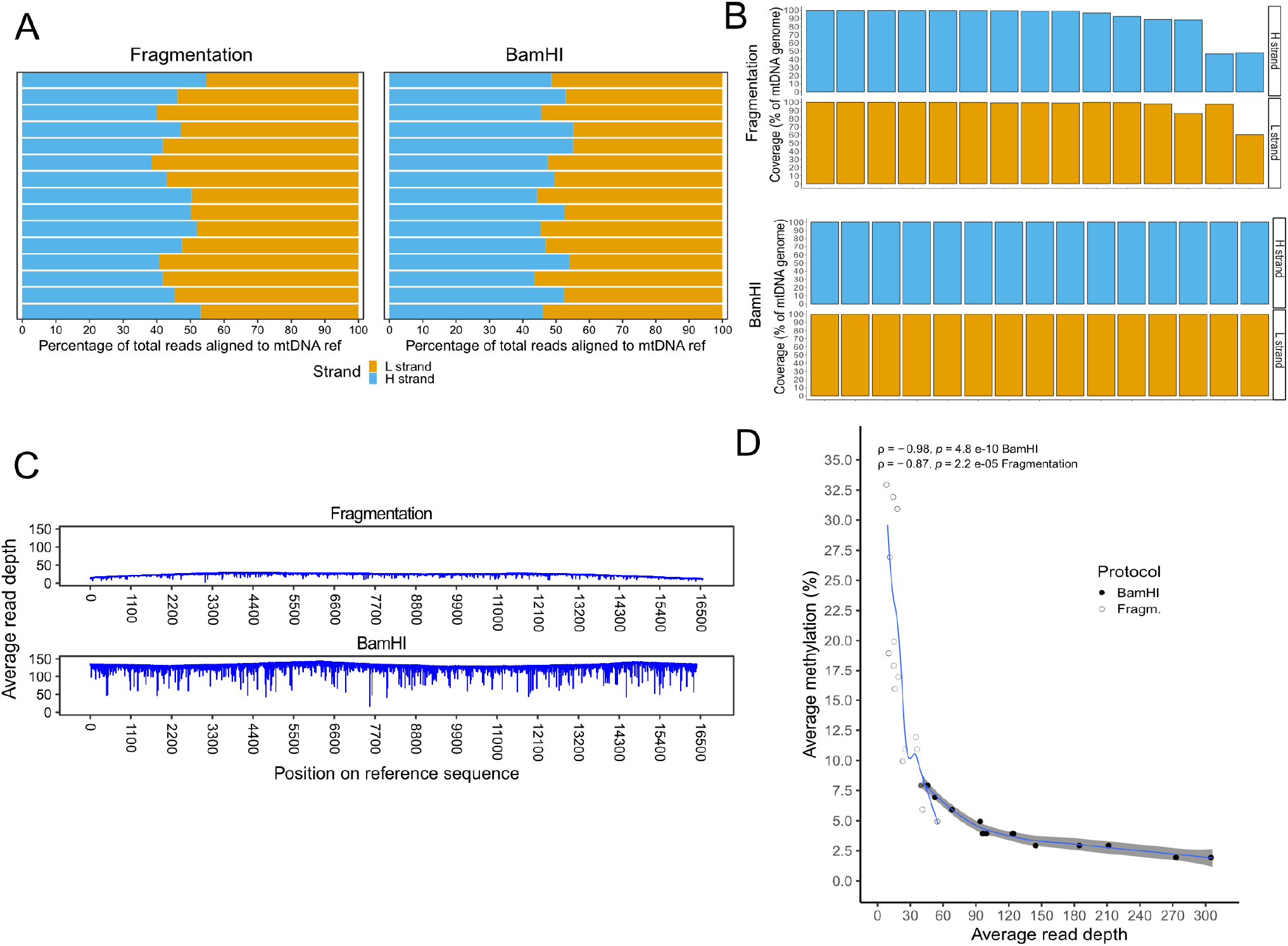
BamHI-based protocol improves mtDNA reads alignment over the standard ONS library preparation. **a)** Percentage of reads aligned to the mtDNA reference per strand and per protocol used (N = 15 samples per protocol). **b)** Percentage of mtDNA covered by at least 5 reads on the two mtDNA strands (H and L) per biological replicate (N = 15 samples per protocol), in samples processed with (top) fragmentation protocol and (bottom) BamHI-based protocol. **c**) Distribution of the average read depth per mtDNA position in samples processed with (top) fragmentation protocol and (bottom) BamHI-based protocol. The red lines indicate the mean over all the data points (calculated using the “loess” *geom_smooth* R function). **d)** Correlation between average read depth and average methylation percentage in samples processed with fragmentation- and BamHI-based protocols. Each dot represents a sample sequenced with either one of the two protocols (N = 15 per protocol). Pearson correlation coefficient and two-sided P-values are shown.

Samples sequenced using a fragmentation-based protocol showed a greater range in average methylation levels (Min_Frag_: 5% − Max_Frag_: 33%) at low read depths levels (Min_Frag_: 9.16x − Max_Frag_: 55.62x) (**Figure 3D**), while BamHI-based protocol samples achieved similar methylation levels (Min_BamHI_: 2% − Min_BamHI_: 8%) at higher read depths (Min_BamHI_: 40.6x − Min_BamHI_: 306.2x) (**Figure 3D**). Overall, these results suggest that our custom-made ONS protocol is more efficient in achieving full-length mtDNA sequences enrichment and higher mtDNA read depths than the standard Nanopore library preparation, with low risk of NUMTs co-amplification.

### ONS-based mtDNA sequencing and CpG methylated sites identification in different human cell lines

Next, we tested the efficacy of the BamHI-based library preparation protocol on 2 groups of human biological samples. First, we sequenced DNA from 3 trans-mitochondrial osteosarcoma cybrids with known mtDNAs belonging to different mtDNA human populations (haplogroups) (Gómez-Durán *et al*., 2012) with an identical nuclear background (Chomyn *et al*., 1994) (Group 1, N = 5 biological replicates of 3 independent DNA from the mitochondrial haplogroup H1, J1c and J2, respectively) (**Data S2**). Then, we sequenced mtDNA from primary fibroblasts including healthy control subjects, and patients carrying heteroplasmic mutation m.8344A>G (*MT-TK)* and m.3243A>G (*MT-TL1*), known to cause mitochondrial respiratory chain enzyme defects at high levels of heteroplasmy (Shoffner *et al*., 1990; Flierl, Reichmann and Seibel, 1997) (Group 2, N = 3 biological replicates, **Data S2**).

First, we applied ONS to detect mtDNA variants and quantify their heteroplasmy. For this, we identified mtDNA variants in ONS-sequenced samples and used high depth Illumina MiSeq sequencing of mtDNA (mean = 2,769x, min = 318x, max = 5,559x) for validation (**Data S3**). Variant calling analysis confirmed ONS haplogroup predictions of Group 1 samples and the detection of the known single nucleotide heteroplasmic variants in Group 2 (**Data S3, Figure S5**). On average, we found 60 mtDNA variants with ≥ 10% heteroplasmy per sample with ONS, of which 28 (∼47%) were confirmed with Illumina Miseq (**Figure 4A** right plot). These were mostly highly heteroplasmic or homoplasmic variants (heteroplasmy_ONS_ = 93% ± 17%; heteroplasmy_Miseq_ = 96% ± 15%; mean ± sd, **Figure 4A** left plot, **Figure S5)**. The remaining unconfirmed mtDNA variants were mostly low heteroplasmic (heteroplasmy_ONS_ = 16% ± 11%, mean ± sd; **Figure 4A** eft plot, **Figure S5**). Heteroplasmies calculated with ONS overall tended to correlate better with Illumina at higher read depths (**Figure S6**). This suggests that adjustments to the ONS protocol aimed at reaching higher read depths (e.g. longer sequencing times, higher starting sample material, etc) can improve heteroplasmic mtDNA variants identification.

**Figure 4.**
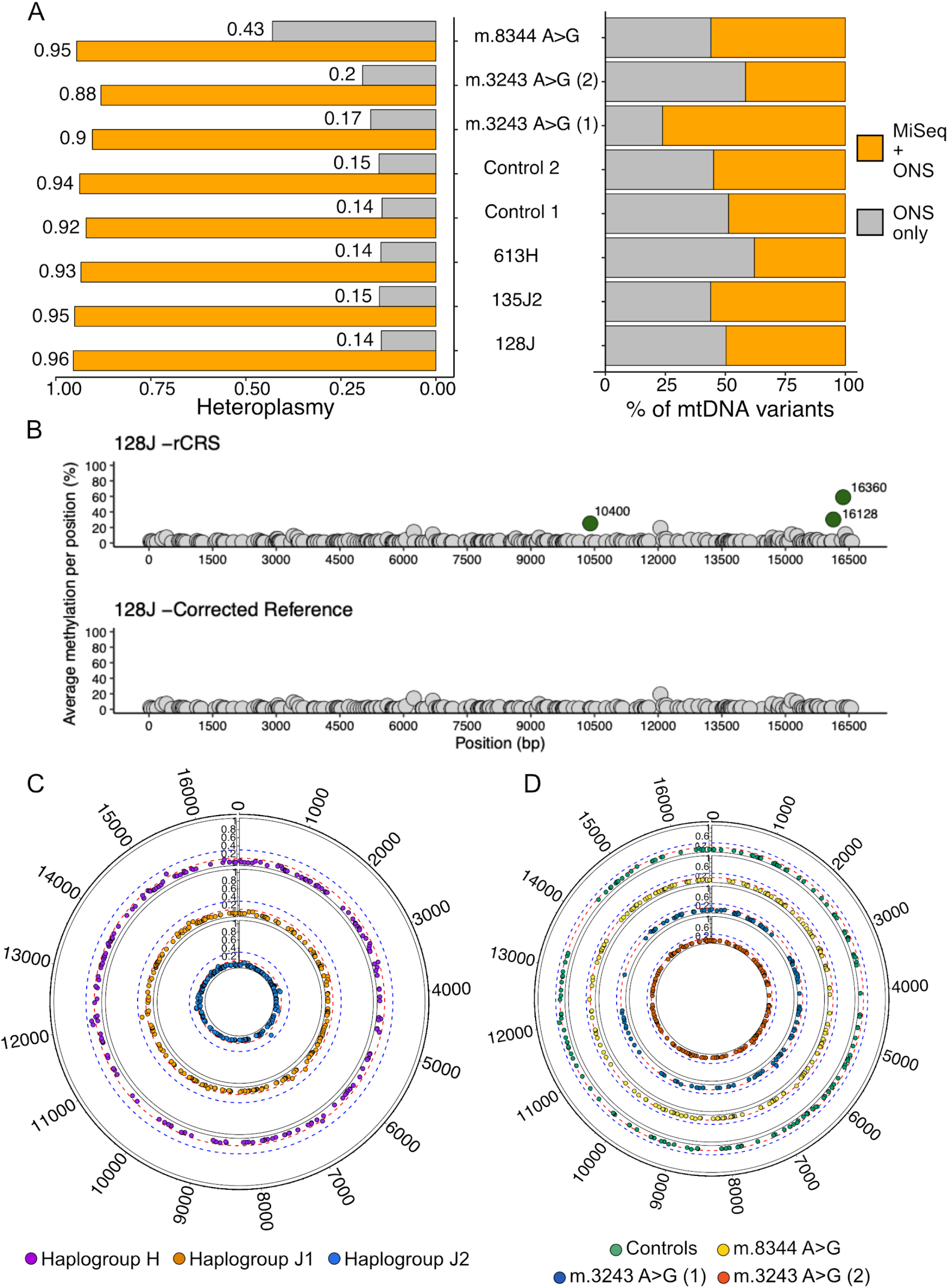
ONS-based variant calling and CpG methylation analysis of mtDNA. **a)** Variant calling statistics per each cell line analyzed. Heteroplasmy (left) and percentage of single nucleotide mtDNA variants (right) identified with either Illumina Miseq and ONS or ONS only. Values are means calculated across all biological replicates per each cell line analyzed. **b)** Example of how average methylation levels change when hg38 (including the mitochondrial reference sequence rCRS) (top) is used versus the sample-specific consensus sequence determined by direct Illumina sequencing (bottom). In green are highlighted the sample-specific differentially methylated positions which disappear upon reference correction. **c-d**) Circos plots showing the CpG methylation levels detected across the whole mitochondrial molecule in Group 1 (**c**) and Group2 (**d**) cell lines. Dashed lines indicate 30% (blue) and 10% (red) methylation levels.

Next, we performed differential methylation (DM) analysis (Gigante *et al*., 2019), which revealed 3 DM-CpGs in Group 1: one found only in the haplogroup J2 (m.16360) cells, and two found in both J cell lines (m.10400 and m.16128) (**Figure 4B** top figure, **Data S4**). We also found 5 DM-CpGs in all Group 2 cells (m.4919, m.9195, m.10400, m.15925, m.16128) (**Data S4**). However, a close scrutiny revealed that an haplogroup-defining variant always fell within a ± 5 bp window from a DM-CpG, prompting us to hypothesize that these variants may alter the Nanopolish methylation calling (Methods). To test this, we generated a new reference for methylation calling based on a mtDNA consensus sequence built on major mtDNA alleles identified with Illumina MiSeq sequencing. DM analysis based on consensus sequence returned no significant differences in methylation levels between the samples, indicating that the previously identified possible DM-CpGs were artefacts of the Nanopolish calling algorithm (**Figure 4B** bottom figure, **Data S4**). Using a sample-specific mtDNA reference sequence for methylation calling, we measured an average of 195 ± 8 (33.3%) methylated CpGs in Group 1 and 141 ± 20 (20%) CpGs in Group 2 with 18 CpG found methylated in both Group 1 and Group 2 samples (**Data S4**). Methylation levels were overall low (**Figure 4C-D**, methylation_GROUP_1_= 0.4%-9.4%, 2%; methylation_GROUP_2_= 0.4%-8.8%, 1.5%; min-max, mean), with only a minority of methylated CpGs shared within groups (27.8% in Group 1 and 9.2% in Group 2; **Figure S7**). Thus, we concluded that neither subtle differences in mitochondrial function linked to population variants (Gómez-Durán *et al*., 2012) nor severe defects caused by heteroplasmic pathological tRNA variants (Shoffner *et al*., 1990; Flierl, Reichmann and Seibel, 1997) differentially affect the levels of mtDNA methylation in human cell lines.

## Discussion

The discovery of mitochondrially-targeted methyltransferases (Shock *et al*., 2011; Wong *et al*., 2013; Patil *et al*., 2019) has prompted the suggestion that mtDNA CpG methylation can be a marker of a variety of diseases including ageing (D”Aquila *et al*., 2015), environmental exposure to tobacco smoke (Vos *et al*., 2020), cancer (Dong, Pu and Cui, 2020) and neurological disease (Blanch *et al*., 2016; Stoccoro *et al*., 2017). Currently, quantitative analysis of CpGs is mostly based either on mass spectrometry or on the bisulfite treatment of gDNA (bisulfite pyrosequencing and WGBS). While the first method is the most sensitive in determining the general CpG methylation level of a given sample, it lacks information about the position of individual methylated residues (Song *et al*., 2005). On the other hand, while bisulfite-based technologies resolve the CpG methylation at a single-base level, they are susceptible to the introduction of biases due to the selective degradation of cytosine-rich sequences (both nuclear and on the L-strand of mtDNA) (Ji *et al*., 2014; Olova *et al*., 2018). Despite this, there is still ample literature which considers WGBS the gold standard technology for the analysis of mtDNA methylation (Dou *et al*., 2019; Patil *et al*., 2019; Sirard, 2019). In an attempt to resolve this controversy, we leveraged 55 publicly available WGBS datasets part of the Roadmap Epigenome Project (Roadmap Epigenomics Consortium *et al*., 2015), focussing on describing per-strand sequencing metrics and how these affect the methylation profile of mtDNA. Our analysis confirmed that in 58% of the samples bisulfite treatment introduces a marked per-strand bias with an impact on global mtDNA CpG methylation levels quantification. In the remaining samples the bias is milder (although present in all of them), influencing local stretches of the mtDNA molecule.

To overcome these limitations, we developed an accurate and reproducible protocol to investigate mtDNA methylation using ONS. We tested our method against the standard ONS library preparation protocol, based on random fragmentation. Our protocol is based instead on selective restriction digestion by BamHI followed by selection of longer sequences, which results in an enrichment for native full-length mtDNA. Also, we demonstrated that selective exclusion of nuclear DNA combined with the *in silico* identification of supplementary alignments, reduces significantly the risk of NUMTs contamination. Comparing our results with Illumina sequencing, we found that our protocol allows the correct calling of the mtDNA nucleotide variants present in the samples, including heteroplasmic pathogenic mutations. Despite this, we also found a large number of unconfirmed low heteroplasmic single nucleotide mtDNA variants, likely due to the error rate intrinsic to ONS chemistry (Dohm et al., 2020). Finally, our analysis also revealed that the methylation calling with Nanopolish is influenced by the presence of mtDNA variants surrounding the CpG residue. In light of this, we recommend a careful review of previously identified methylated positions and of differential methylation results (Goldsmith *et al*., 2020).

Overall, our study indicates that, after removing the technical biases intrinsic to previously used techniques and to ONS, there is negligible mtDNA methylation in normal or cancer cell lines, at least in the conditions analysed. In conclusion, we recommend the use of our protocol for the analysis of epigenetic modifications of mtDNA at single-molecule level and that WGBS-based studies that show evidence of mtDNA methylation should be carefully re-considered (Dou *et al*., 2019; Patil *et al*., 2019; Sirard, 2019).

### Limitations of Study

To reproduce this protocol, we suggest starting with at least 2µg of DNA before the BamHI digestion. BamHI was specifically chosen because it usually cuts human DNA only once. Different restriction enzymes will be needed when extending this approach to other species. This protocol may be used to study different kinds of mtDNA epigenetic modifications (such as 6mA) at a single-molecule and nucleotide base resolution.

## Acknowledgments

The authors thank Dr. Lisa Tilokani, Dr. Michele Frison and Dr. Robert Preston for their excellent scientific and technical assistance. I.B. is funded by the Cambridge Trust and the MRC-DTP PhD Program. A.G.D is an Atracción de Talento M1 Fellow from Comunidad de Madrid (Spain) (2019-T1BMD-14236). P.F.C. is a Wellcome Trust Principal Research Fellow (212219/Z/18/Z), and a UK NIHR Senior Investigator, who receives support from the Medical Research Council Mitochondrial Biology Unit (MC_UU_00015/9), the Evelyn Trust, and the National Institute for Health Research (NIHR) Biomedical Research Centre based at Cambridge University Hospitals NHS Foundation Trust and the University of Cambridge.

## Author contributions

I.B., A.G.D., C.C and P.F.C designed the study; A.G.D., C.C and P.F.C oversaw the study; I.B., performed the ONS sequencing and experiments, Z.G. performed the Illumina sequencing; I.B. and C.C. analysed the data; I.B., C.C., A.G.D. and P.F.C. wrote the manuscript. The authors read and approved the final manuscript.

## Data Legends

**Data S1: List and metrics of WGBS samples that passed quality control**

The table includes for each bias group (tabs “Bias” and “Low Bias”): sample IDs and descriptions, total sequenced throughput (in basepairs), average mitochondrial read depths measured with Samtools depth, number and percentage of reads aligned to each mitochondrial strand, coverage per strand.

**Data S2. List and metrics of samples sequenced with ONS in this study**.

The table includes: sample IDs and descriptions, library preparation method used for sequencing with ONS (fragmentation/BamHI), total sequenced throughput (in basepairs), average mitochondrial read depths measured with Samtools depth, number and percentage of reads aligned to each mitochondrial strand, coverage per strand.

**Data S3. Illumina Miseq and ONS sequencing metrics and variant calling**

The table includes: Miseq and ONS sequencing read depth and coverage (percentage of mtDNA covered by at least one read) calculated by running the MToolBox pipeline (tab “Read depth, coverage, haplo predictions”); haplogroup predictions calculated with MToolBox and Haplogrep2 (tab “Read depth,coverage, haplo predictions”); list of mtDNA SNVs identified with ONS, with read depth ≥ 30 and variant allele fraction ≥ 10% and corresponding *per*-base read depth and allele fraction quantified with Illumina Miseq (tab “mtDNA variants”).

**Data S4: ONS methylation analysis results**

The table includes: average methylation values per position calculated on Group1 and Group2 samples using the consensus sequence calculated on Illumina results (tabs “Group1 meth. values”“ and “Group2 meth. values.”). The 18 methylated CpGs shared by the two groups are highlighted in bold; differential methylation analysis results performed on average methylation values calculated with rCRS (tab “Diff. meth. on rCRS) or the Illumina consensus sequence (tab “Diff. meth. on consensus).

## Methods

### Cell culture

Cell lines used in this study are listed in the Key Resources Table. Cells were maintained in fibroblast medium [DMEM high glucose (Gibco) with 10% fetal bovine serum (Gibco) and no antibiotics] at 37°C in a humidified 5% CO2 atmosphere. Cells were grown until ∼80% confluence. When ready, cells were washed with PBS (Gibco), then incubated with 0.05% trypsin (Gibco) for 5 minutes at 37°C. Cells were collected by centrifugation (1500 rcf for 5 minutes) and pellets were washed once with PBS, before being snap frozen in liquid nitrogen and kept at −20°C until further use.

### DNA extraction and DNA quantification

All DNA used in this study was extracted from snap-frozen pellets using the QIAmp DNA mini kit (QIAGEN) following the manufacturer”s instructions. DNA was quantified using the Qubit dsDNA kit (Invitrogen) following the manufacturer”s instructions.

### Long-range polymerase reactions (LR-PCR)

LR-PCR amplification reaction was performed using PrimeSTAR GXL DNA Polymerase kit (Takara) according to manufacturer”s instructions. The primers used are detailed in **Table S2**. Product length encompasses most part of the mtDNA sequence. Amplification reactions were performed using the following cycling conditions: 94°C for 1 minute, followed by 30 cycles of 98°C for 10 seconds, 55°C for 15 seconds and 68°C for 10 minutes.

### Generation of negative and positive controls

Untreated LR-PCR amplicons were used as negative controls for methylation. To generate positive controls, the same amplicons were treated *in vitro* with the recombinant CpG methyltransferase M.SssI (NEB). Briefly, 1µg of amplicon DNA per 50µl reaction was treated for 4 hours at 37°C with 50 units of M.SssI in the presence of 1x NEB buffer #2 and 160µM of S-adenosylmethionine (SAM). To test the efficiency of the M.SssI reaction, 10 units of methylation-sensitive restriction enzyme BstUI were added at the end of the incubation. This was followed by a further incubation at 60°C for 1 hour.

Protection of the M.SssI-treated amplicons from BstUI digestion was assessed using the Genomic DNA ScreenTape System (Agilent) on an Agilent 2200 TapeStation platform following manufacturer”s instructions (data not shown).

### Mitochondrial DNA enrichment for single-molecule sequencing

1 µg of genomic DNA (nuclear + mitochondrial DNA) per 50 µl reactions was digested with 40 units of the recombinant restriction enzyme BamHI-HF (NEB) for 1 hour at 37°C in the presence of CutSmart buffer (NEB). To achieve combined DNA purification and selection of high molecular weight fragments, DNA was purified using Monarch® PCR & DNA Cleanup Kit (NEB), using the following recommended protocol modification: 20µl of elution buffer was heated to 50°C before the last elution step.

### Quantification of mtDNA levels using ddPCR

ddPCR was used to quantify relative mtDNA enrichment following BamHI-HF (NEB) treatment of control DNA. To quantify relative mtDNA copy number (Krueger and Andrews, 2011), a mitochondrial and nuclear target (the genes *MT-ND1* and *RNASE P*, respectively) were amplified and fluorescent signal was generated using the primers and probes detailed in the **Table S2**. ddPCR protocol was performed following manufacturer”s instructions. Briefly, PCR reaction master mix was prepared in 1x (final concentration) ddPCR Supermix for Probes (No dUTP, BioRad), by adding 300nM of each primer and 200nM of each probe in 19µl final volume. 1 ng of sample DNA was then added to the mastermix. Droplets were generated using an Automated Droplet Generation instrument (BioRad) and were then subjected to PCR amplification, performed using the following cycling conditions: 95°C for 10 minutes, followed by 39 cycles of 94°C for 30 seconds and 58°C for 1 minute, followed by a final stabilisation step at 98°C for 10 minutes. Droplets were then loaded into a QX200 droplet reader (BioRad) and analysed using an absolute quantification protocol (ABS) to measure the absolute copy number of each probe. Droplet analysis was performed using the QuantaSoft analysis software (BioRad).

### ONS library preparation and sequencing on the MinION instrument

Approximately 1 µg of native genomic DNA or purified LR-PCR amplicons were prepared for ONS sequencing on R9.4.1 flow cells using the Ligation Sequencing Kit SQK-LSK109 (Nanoporetech), in combination with the Native Barcoding Expansion Kit EXP-NBD114 (Nanoporetech). Genomic DNA was fragmented either through BamHI digestion (Methods) or sheared to 10 kbp using g-tubes (Covaris), following manufacturers” instructions. Simultaneous DNA repairing, end-repairing and dA-tailing was achieved using the NEBNext FFPE Repair Mix (NEB) and the Ultra II end-repair module (NEB). Barcodes were ligated to individual samples using Blunt/TA Ligase Master Mix (NEB). Samples were then combined and AMII adapters containing the motor proteins needed for sequencing were ligated using NEBNext® Quick Ligation Module (NEB). AMPure XP beads (Beckman Coulter) at a concentration of 1x, 1x and 0.5x, respectively, were used to purify DNA between the library preparation steps. Final libraries were loaded onto R9.4.1 flow cells and samples were sequenced using a single MinION Mk 1B. To keep the sequencing throughput consistent, 6 biological samples were always pooled together and sequenced for 24 hours. LR-PCR amplicons were pooled together and sequenced for 6 hours.

### Illumina Miseq library preparation and sequencing

MiSeq libraries were prepared from genomic DNA by amplification of the mitochondrial DNA in two overlapping fragments (Weissensteiner *et al*., 2016), using the primers outlined in the Key Resources Table. Amplicons were individually purified, quantified, and then were pooled in equal amounts from each sample. Libraries were prepared using NEBNext Ultra library prep reagents (NEB) according to manufacturer”s instructions and sequenced using a 2 × 250-cycle MiSeq Reagent kit v3.0 (Illumina, CA).

### WGBS data analysis

Raw WGBS experiments part of the Roadmap Epigenome Project (Roadmap Epigenomics Consortium *et al*., 2015) were downloaded from the GEO Database. Downloaded files from single-ended WGBS sequencing experiments were converted from SRA format to fastq files using `fastq-dump` (Key Resources Table) with the following options: `╌readids` ╌`skip-technical −W ╌read-filter pass ╌gzip`. Read quality of the converted fastq files was assessed with FastQC v0.11.5 (Andrews, 2015). All of the reports generated from FastQC were manually checked to determine whether a trimming of low-quality reads and/or adapters was needed. Where trimming was deemed necessary, TrimGalore! v0.4.5 (Krueger, 2016) was used. The software automatically trims adapter sequences from the reads (if present) and retains those with an average Phred quality score ≤ 20 (before and/or after trimming). Reads shorter than 45 bp after trimming were discarded using `╌length` option. Upon quality check and trimming, both alignment of the WGBS fastq files to the reference human genome sequence (GRCh38) and extraction of the methylation information were carried out with bowtie2 v2.3.2 (Langmead and Salzberg, 2012) and Bismark v0.19.0 (Krueger and Andrews, 2011), respectively. Coverage was calculated from BAM files using `samtools depth`. This was defined as the percentage of mtDNA genome in each strand covered by at least 5 reads. Methylation extraction was carried out using the `bismark_methylation_extractor` package with the following options: `╌comprehensive ╌merge_` `non_CpG ╌gzip ╌ bedGraph ╌CX_context`, but only CpG residues were considered for further analyses.

### ONS data analysis

Base-calling of fast5 files containing raw electric current information was performed by the `guppy_basecaller` package of Guppy v3.2.2+9fe0a78 (Nanoporetech). Base-called, barcoded reads were de-multiplexed into individual samples using the `guppy_barcoder` package of Guppy v3.2.2+9fe0a78 (Nanoporetech). In order to simultaneously enrich for linear full-length mitochondrial sequences, exclude ligation artifacts and minimise the presence of NUMTs, we applied a stringent filter on read sequence length (min: 4000 bp, max: 17000 bp) and quality (Phred quality score ≥ 9) using NanoFilt v2.2.0 (De Coster *et al*., 2018) on the barcoded fastq files (**Figure 2F**).

Minimap2 v2.10-r761 (Li, 2018) with the `-x map-ont` option was used to perform the alignment of Nanopore reads onto the GRCh38 reference (which includes the mitochondrial rCRS reference sequence, NC_012920.1), and the option `-secondary=no` was used to exclude secondary alignments in the BAM output. Because minimap2 does not recognise circular reference sequences, reads spanning the D-loop are reported as supplementary alignments in the output BAM files. For this reason, we included in the final set of aligned reads also supplementary alignments aligning onto the mtDNA reference and spanning the D-loop, but only if they aligned in the same orientation on the same strand (H or L strand). Any other kind of supplementary alignment was excluded. Similarly, to avoid the same issue with reads spanning the BamHI cut site in the *ND6* gene (base 14258-14259 of the mtDNA reference sequence), we created an alternative GRCh38 reference sequence with a modified mitochondrial reference starting at base 14259 instead of base 1. All of the experiments where the samples were fragmented using BamHI were aligned to this alternative sequence (gene annotations were adapted accordingly). Quality control plots and sequencing statistics were automatically generated using NanoPlot v1.13.0 (De Coster *et al*., 2018).

### ROC curve generation

We calculated a ROC curve to assess the accuracy of our methylation calling, using a procedure previously adopted in (Simpson *et al*., 2017). Briefly, we randomly chose 50,000 mtDNA CpG sites from positive and negative controls and classified each CpG call as true positive (TP) or false positive (FP), depending on which of the two controls each site came from and on whether methylation fell above or below a log-likelihood methylation threshold. We repeated the TP and FP calculation by varying log-likelihood threshold values within a range of −20 to 20 (to build the ROC curve) and 0 to 10 (to calculate accuracy, intended as the proportion of true calls, either TP or true negatives (TN)), with a step of 0.25, as explained in (Simpson *et al*., 2017).

### Mitochondrial variant calling of ONS samples

Because Nanopore technology allows a simultaneous read of epigenetic modifications while sequencing the target DNA, we performed a mitochondrial variant calling on the fastq files filtered with NanoFilt v2.2.0 (De Coster *et al*., 2018). For this we used a modified version of the MToolBox pipeline (Calabrese *et al*., 2014), adapted to long-reads sequencing analysis (Key Resources Table). Briefly, the main changes integrated into the MToolBox workflow are 1) the use of the Minimap2 aligner software (Li, 2018) for long-reads mapping and 2) additional parsing of SAM files to include reads uniquely mapped on the mtDNA reference and reads with supplementary alignments but only showing mtDNA mapping locations. These reads can be the results of the process of linearization of the circular molecule of mtDNA due to random fragmentation or to BamHI enzymatic cut. Reads with secondary or supplementary alignments on the nuclear genome were excluded and classified as possible NUMTs. For read mapping we used the GRCh38 human genome assembly (which includes rCRS as mitochondrial reference sequence). For variant calling, we set the quality score (QS) threshold to retain variants to 10 (changing the `-q` option of the `assemblyMTgenome.py` script). Variants with a read depth per position ≥ 30 and variant allele fraction ≥ 10% were retained. Only single nucleotide variants (SNVs) were considered for comparison with Illumina Miseq sequencing. Haplogroup predictions were performed using both MToolBox and Haplogrep 2 v.2.1.1 (Weissensteiner *et al*., 2016). Haplogrep2 predictions were based on homoplasmic variants only (with variant allele fraction ≥ 0.9).

### Mitochondrial variant calling of Illumina Miseq samples

Fastq files generated with Illumina Miseq were checked for quality using FastQC v0.11.5 (Andrews, 2015). Illumina adapters and read ends showing poor *per*-base quality were trimmed using TrimGalore! v0.4.5 (Krueger, 2016), setting a minimum per-base QS = 20 and minimum read length after trimming = 35 bp. Mitochondrial variant calling was then performed with the standard MToolBox pipeline (Calabrese *et al*., 2014), which mapped reads to the human reference genome (GRCh38) with the two-mapping step protocol, to exclude possible NUMT. Single nucleotide variants with ≥ 5 reads of support (and at least 1 read of support on each strand) and minimum QS per base ≥25 were retained. Haplogroup predictions were performed using both MToolBox and Haplogrep 2 v.2.1.1 (Weissensteiner *et al*., 2016). Haplogrep2 predictions were based on homoplasmic variants only (with variant allele fraction ≥ 0.9).

### CpG methylation detection in ONS samples

Detection of methylation in CpG context was carried out using Nanopolish v0.11.0 `call-methylation` package (Simpson *et al*., 2017). Nanopolish utilises a trained Hidden Markov Model to detect modified bases by comparing raw electric signals of modified/unmodified cytosines with expected signal from a reference sequence. The methylation calling output is a log-likelihood ratio where a positive value indicates evidence supporting methylation. Nanopolish utilises `fast5` files containing raw electric signal information, basecalled fastq files and BAM alignment files to generate an index file used by the algorithm to determine methylation Log-likelihood ratios. Minimap2 alignments to reference sequences were performed with the same parameters described in the Nanopore Data Analysis section. Log-likelihood ratios were then converted to a binary methylated/unmethylated call for each read, then percentage of methylation was obtained by calculating the fraction of methylated reads, using the `calculate_methylation_frequency.py` script available in the package. The default calling threshold of ≥ 2.5 LLR was modified to a more stringent ≥ 5 LLR to increase the accuracy of the call. Since Nanopolish groups neighbouring CpG sites and calls them jointly, CpG sites in the same group were separated and assigned the same methylation frequency using the `-s` option.

### CpG methylation analysis in ONS samples

We applied a series of stringent quality filters to remove possible artefacts of the CpG methylation calling and errors introduced by the Nanopolish algorithm. We first removed CpGs calls with a methylation frequency greater than two standard deviations from the mean in negative controls (false positives, **Table S1**). We also removed: i) calls supported by less than 60 reads; ii) calls with methylation frequency similar to the background (i.e. ≤ 0.5%, the average methylation frequency observed in the negative control) and iii) calls neighbouring any heteroplasmic nucleotide variant (heteroplasmy < 0.9) in a ± 5 nucleotides window. This last approach was deemed necessary after noticing that Nanopolish introduced a false methylation call every time a homoplasmic haplogroup-defining variant position fell within ± 5 nucleotides from a CpG. As 11 nucleotides is the kmer size that Nanopolish considers to calculate CpG LLR, we hypothesized that the introduction of a nucleotide variant within ± 5 nucleotides from the CpG altered the Nanopolish methylation determination, leading to an incorrect methylation call. To demonstrate this, we used MToolBox (Calabrese *et al*., 2014) to generate a consensus sequence from the Illumina data, carrying the major alleles at each position, and used this new sequence to perform the methylation calling again on ONS samples. As expected, this time no methylation was identified in the CpGs close to the haplogroup-defining variants (**Figure 4B**). Differential methylation analysis was performed using the R package DSS (Park and Wu, 2016), following the protocol detailed in (Gigante *et al*., 2019) using the H haplogroup and control fibroblasts as baseline in Group 1 and Group2, respectively. Differentially methylated mtDNA positions and regions (defined by overlapping tiles of 50nt) were deemed significant if False Discovery Rate was below 1%.

## Statistical test

Unless stated otherwise, pairwise comparisons were tested for significance using Wilcoxon two-tailed test.

## Data and code availability

Raw sequencing data will be made available upon publication on the SRA archive. The MToolBox pipeline used for mitochondrial variant calling and code for filtering VCF files are available as GitHub branch of the MToolBox repository: https://github.com/mitoNGS/MToolBox/tree/MToolBox_Nanopore. Code used for plotting and data analysis is available at https://github.com/ib361/scripts_paper.

