## Supplemental information for "Oxford Nanopore sequencing-based protocol to detect CpG methylation in human mitochondrial DNA"

### Supplemental Figures

#### Figure S1.WGBS alignment bias and methylation analysis, related to Figure 1

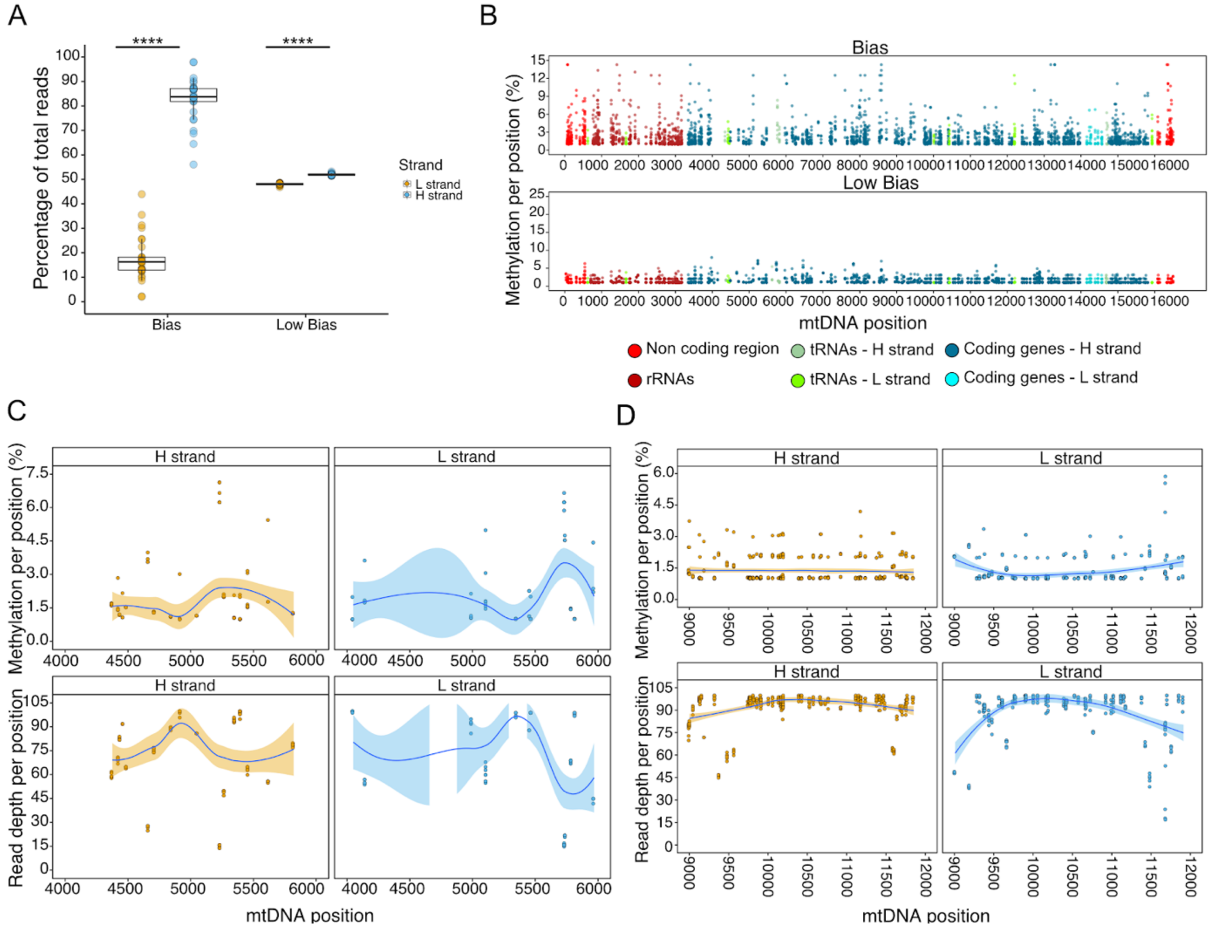


**a)** Percentage of reads aligned to mtDNA, divided by bias group. Boxplot shows the percentage of reads aligned to the mtDNA reference. The lower and upper hinges correspond to the first and third quartile of the distribution, with median in the center and whiskers span no further than 1.5*interquartile range. Stars indicate significance (****: two-sided P ≤ 0.0001, Wilcoxon test). **b**) Distribution of the methylation percentage per mtDNA position in CpG context, in (top) Bias (N = 32) and (bottom) Low Bias (N = 23) groups.  Each dot represents a sample. Methylation values are expressed in % of methylation. **c-d**) CpG methylation and read depth profiles of a 2kb (**c**) and 3kb (**d**) mtDNA genome region, per each position in the Low Bias sample group, divided by mtDNA strand (H and L). Each dot represents a sample. Methylation values are expressed in % of methylation. Blue lines indicate the mean over all the data points (calculated using the "loess" *geom_smooth* R function) and shaded surrounding regions represent 95% confidence interval.

**Figure S2. Experimental setup of mitochondrial negative and positive controls and ONS methylation analysis, related to Figure 2**

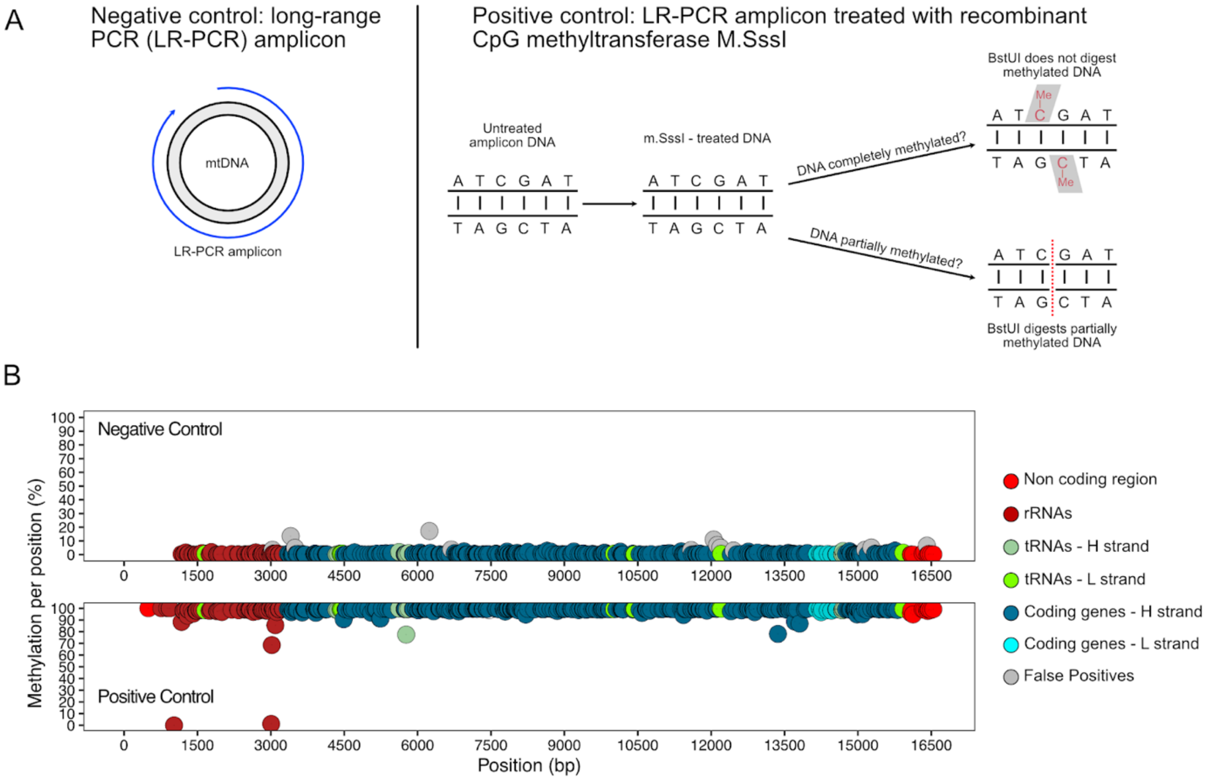


**a**) Experimental setup for the generation of negative and positive controls. **b**) Distribution of the methylation percentage per mtDNA position in CpG context, in (top) negative and (bottom) positive controls. Methylation values are expressed in % of methylation.

**Figure S3. Nanopore sequencing metrics**

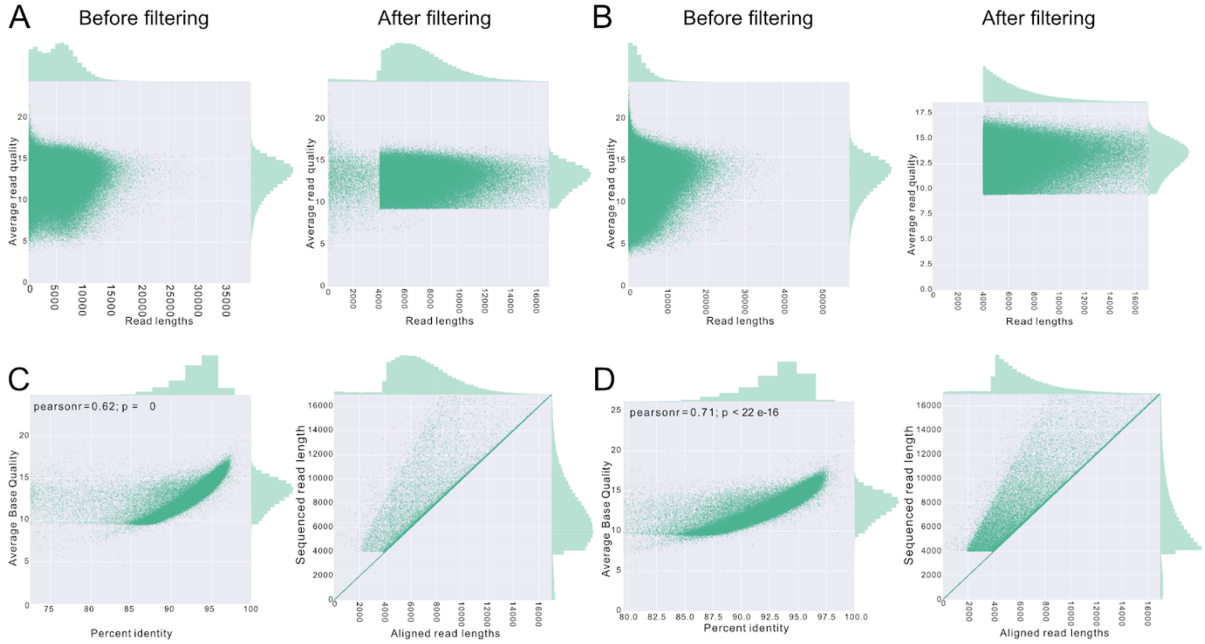


**a)** Plots show the correlation between read lengths and read quality scores in one sample processed with the fragmentation protocol before filtering (left) and after filtering (right). **b)** Plots show the correlation between read lengths and read quality scores in one sample processed with the BamHI-based protocol before filtering (left) and after filtering (right). **c)** Plots show the correlation in one sample processed with the fragmentation protocol between percent identity to the reference sequence and average quality of the reads (left), and correlation between aligned read lengths and sequenced read lengths (right). **d)** Plots show the correlation in one sample processed with the BamHI protocol between percent identity to the reference sequence and average quality of the reads (left), and correlation between aligned read lengths and sequenced read lengths (right).

**Figure S4. ONS alignment metrics, related to Figure 3**

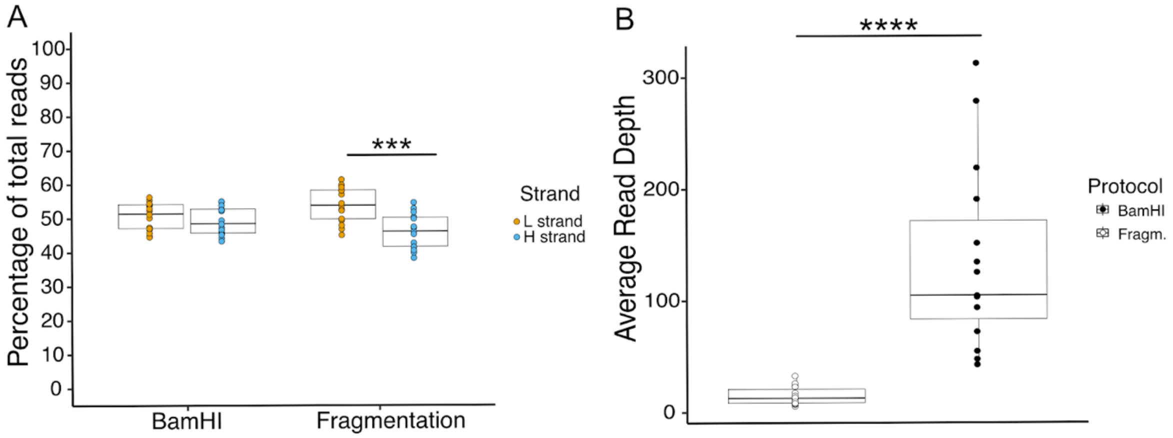


**a)** Percentage of reads aligned to mtDNA, divided by mtDNA strand and library preparation protocol (N = 15 per protocol). Stars indicate significance (***: P ≤ 0.001, Anova one-way test). **b)** Average read depth per sample observed in the same sample pool processed using either fragmentation protocol (left) and BamHI-based protocol (right). N = 15 per protocol. Stars indicate significance (****: two-sided P = ≤ 0.0001, Wilcoxon test).

**Figure S5. mtDNA variants identified with ONS, related to Figure 4**

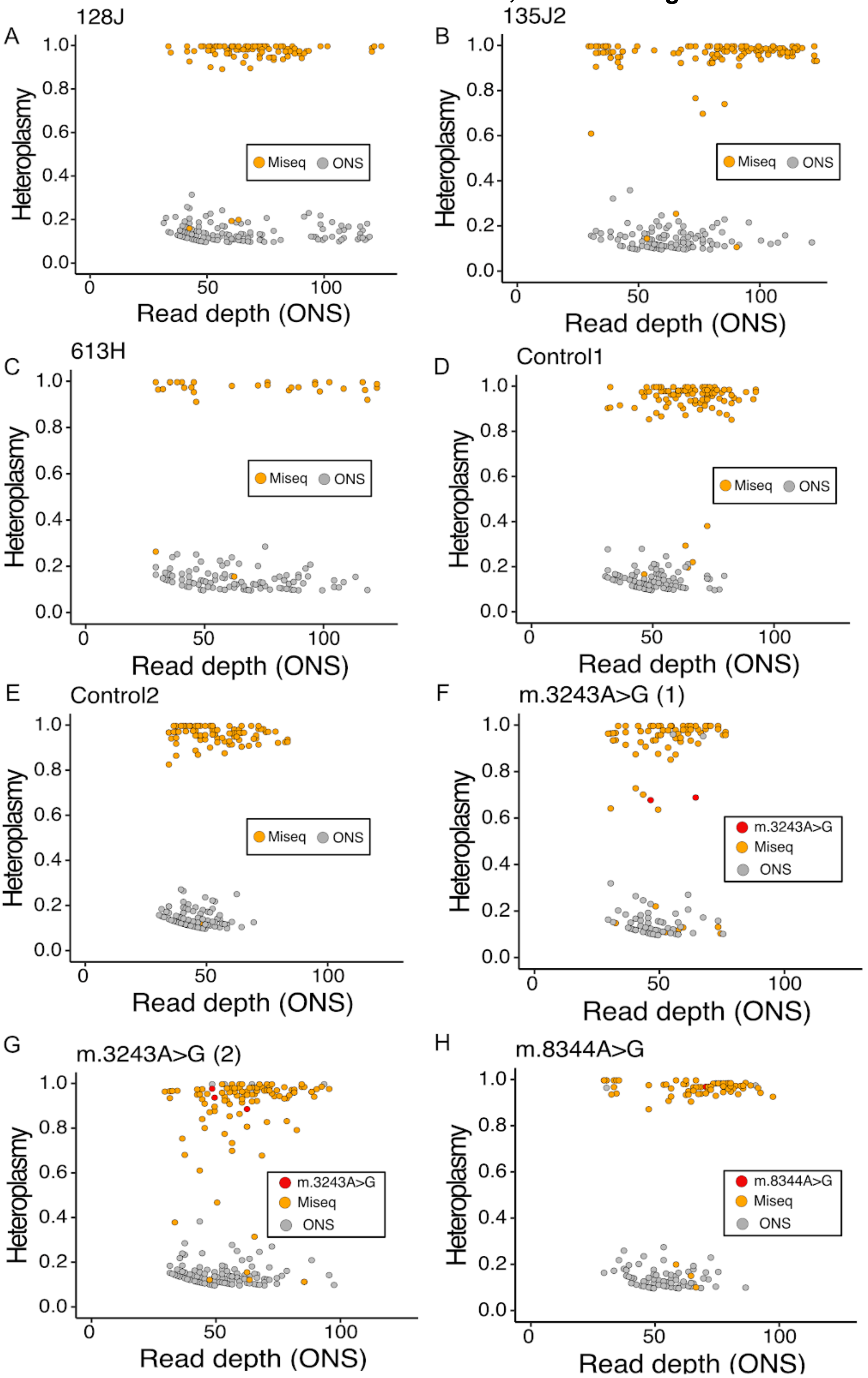

**a**-**h**) Scatterplots show the mtDNA heteroplasmy quantified with ONS in each sample as a function of the read depth per position. Variants confirmed also by Illumina Miseq are highlighted in orange and in red (marking m.3243A>G and m.8344A>G mtDNA mutations). MtDNA variants shown have been aggregated across biological replicates per sample sequenced with ONS (N = 5 for 613H/128J/135J2 and N = 3 for Control1/Control2/m.3243A>G (1)/m.3243A>G (2)/m.8344A>G). The m.3243A>G mutation was confirmed by Illumina Miseq but identified with ONS in two out of three biological replicates of the m.3243A>G (1) sample and in all the three replicates of the m.3243A>G (2) sample. The m.8344A>G mutation was confirmed by Illumina Miseq and identified by ONS in two out of the three biological replicates of the m.8344A>G sample.

**Figure S6. Differences in heteroplasmy detection as a function of the ONS read depth, related to Figure 4**

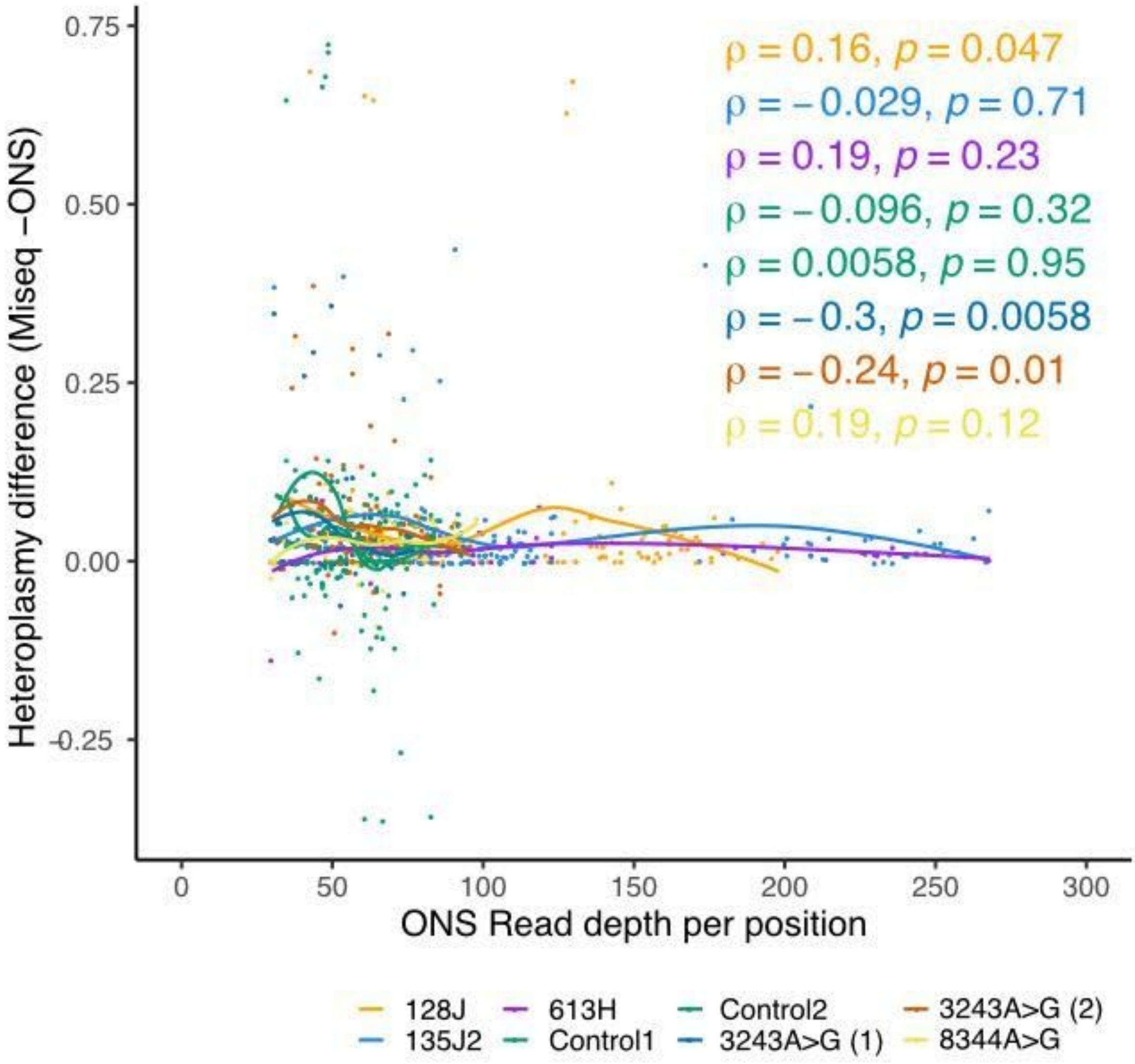


Scatterplot showing a correlation between differences in heteroplasmy values quantified with Illumina Miseq and ONS (calculated as Miseq heteroplasmy - ONS heteroplasmy), for each single nucleotide mtDNA variant detected with both techniques, and ONS read depth per position. Colors correspond to the different samples analyzed (N = 5 for 613H/128J/135J2 and N = 3 for Control1/Control2/m.3243A>G (1)/m.3243A>G (2)/m.8344A>G), with lines indicating mean over all the data points in each sample (calculated using the “loess” *geom_smooth* R function). Spearman's rank two-sided P-values and rho coefficients are shown.

**Figure S7. Sample-specific and shared CpG methylated sites identified with ONS**

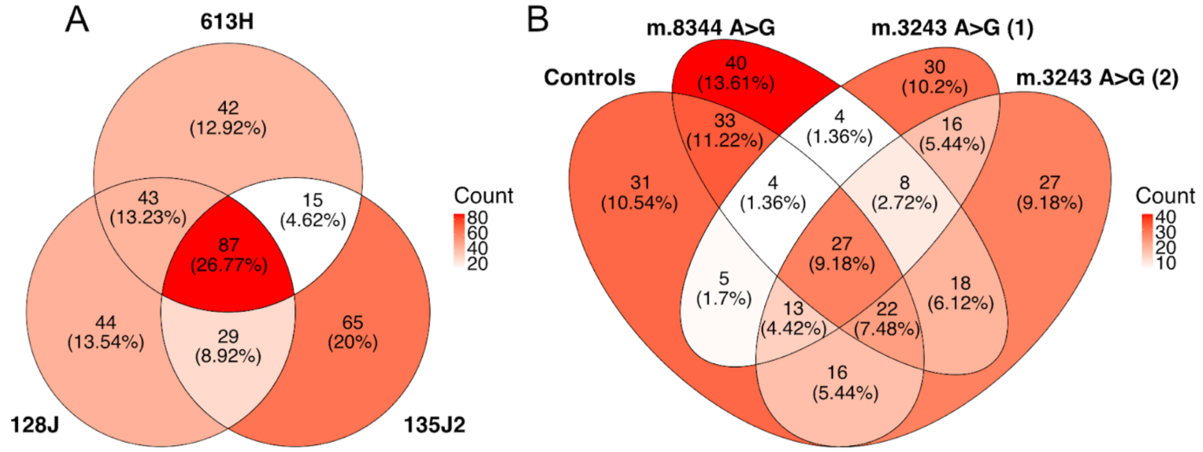


**a-b**) Venn diagrams showing shared and sample-specific methylated CpGs considering all methylated positions found in the mtDNA. Absolute numbers and percentages calculated across all methylated positions in each set are shown.

### Supplemental Tables

#### Table S1. False positives position list

| MtDNA position | Methylation Frequency |
|---|---|
| 3034 | 0.035 |
| 3405 | 0.135 |
| 3494 | 0.05 |
| 6241 | 0.172 |
| 6688 | 0.037 |
| 11590 | 0.033 |
| 12052 | 0.11 |
| 12123 | 0.07 |
| 12190 | 0.051 |
| 12455 | 0.03 |
| 15146 | 0.035 |
| 15274 | 0.052 |
| 16410 | 0.065 |

### Table S2. Primers and probes list

| **Primers** | | | |
|---|---|---|---|
| Primer name | Forward 5' - 3' | Reverse 5' - 3' | Used for |
| 2F | TGTAAAACGACGGCCAGTTTAAAACTCAAAGGACCTGGC | - | LR-PCR |
| D1R | - | CAGGAAACAGCTATGACCAGGGTGATAGACCTGTGATC | LR-PCR |
| MT-ND1 | GGGTTCATAGTAGAAGAGCGATGG | ACGCCATAAAACTCTTCACCAAAG | dPCR |
| RNASE P | AGATTTGGACCTGCGAGCG | GAGCGGCTGTCTCCACAAGT | dPCR |
| **Probes** | | | |
| Probe name | Fluorophore | Sequence 5' - 3' | Quencher |
| MT-ND1 | HEX | ACCCGCCACATCTACCATCACCCTC | BHQ_1 |
| RNASE P | FAM | TTCTGACCTGAAGGCTCTGCGCG | BHQ_1 |
